# Inducible Yeast Two-Hybrid with Quantitative Measures

**DOI:** 10.1101/2021.07.01.450807

**Authors:** Jesus Hernandez, Kevin D. Ross, Bruce A. Hamilton

## Abstract

The yeast two-hybrid (Y2H) assay has long been used to identify new protein-protein interaction pairs and to compare relative interaction strengths. Traditional Y2H formats may be limited, however, by use of constitutive strong promoters if expressed proteins have toxic effects or post-transcriptional expression differences in yeast among a comparison group. As a step toward more quantitative Y2H assays, we modified a common vector to use an inducible *CUP1* promoter, which showed quantitative induction of several “bait” proteins with increasing copper concentration. Using mouse Nxf1 (homologous to yeast Mex67p) as a model bait, copper titration achieved levels that bracket levels obtained with the constitutive *ADH1* promoter. Using a liquid growth assay for an auxotrophic reporter in multiwell plates allowed log-phase growth rate to be used as a measure of interaction strength. These data demonstrate the potential for quantitative comparisons of protein-protein interactions using the Y2H system.

## INTRODUCTION

Identifying quantitative changes in protein-protein interaction (PPI) networks caused by allelic variation in a protein of interest remains technically challenging. This is especially true for natural variation consistent with grossly normal protein function, where the expectation might be difference in relative strength of interactions rather than qualitative differences in number or identity of interactions. Potential for covariation among protein sequence, abundance, and conformational states further complicates most simple assays. An additional complexity for cellular assays, which may provide needed context for some interactions, is that high expression of the tested protein interfaces may by itself pose toxicity, complicating the interpretation of terminal readout measures. The mRNA nuclear export factor NXF1 may serve as an example: homologous to yeast MEX67p, it is an essential gene with a highly conserved core PPI network [1–11], yet a single amino acid substitution in mice (E610G) creates a potent genetic suppressor of intronic retroviral insertion mutations without changing apparent steady-state NXF1 protein levels [12–14].

One approach to identifying quantitative differences in interaction strength is to test the strength of reporter expression in a yeast two-hybrid (Y2H) assay [15, 16]. This easily-modified approach has several variants that have in common use of a hybrid protein with a sequence-specific DNA binding domain (e.g., GAL4-DBD) fused to a protein of interest (the “bait”) and a second hybrid protein with a transcriptional activation domain fused to a potential bait-interacting protein (the “prey”). Physical interaction between bait and prey in the nucleus constitutes a bipartite transcription factor that drives expression of one or more reporter genes in proportion to the level of expression and strength of interaction between the bait and prey. Prior work has used both colonial growth on solid media and liquid growth combined with auxotrophic, enzymatic, or fluorescent reporters [15–21], but typically do not allow adjustment for differences in expressed protein levels.

Using a strong constitutive promoter to drive high-level expression generally provides good sensitivity for interactions, but can be a limitation in some applications. The classic GAL4-based Y2H plasmids, for example, use a constitutive *ADH1* promoter. In studies preliminary to the work presented here, application of this system to allelic Nxf1 bait proteins from divergent mouse strains, C57BL/6J (B6) and CAST/EiJ (CAST), achieved relatively low and unequal expression levels from the *ADH1* promoter in yeast, potentially by competing with endogenous Mex67p for a more limiting factor. To address this, we modified the pGBKT7 bait vector to include a synthetic version of the copper-inducible *CUP1* promoter, which can be quantitatively induced by addition of copper ions to the media [22–26]. The modified vector allowed titratable expression of multiple fusion proteins and allowed functional titration of Y2H interaction using growth rate in liquid culture as an assay for an auxotrophic reporter. Varying copper ion concentration from 0 to 200 μM permitted a range of expression levels that bracketed the level achieved with the *ADH1* vector. Inducible Y2H should have applications for bait proteins that are mildly toxic under constitutive conditions or for quantitative comparisons of affinity between bait proteins with different steady-state levels in yeast, by allowing for interaction measures across expression levels.

## MATERIALS AND METHODS

### Plasmid construction

Yeast Matchmaker Two-Hybrid System was obtained from Clontech (now Takara Bio), including pGBKT7 bait vector, pGADT7 prey vector, and pGBKT7-*Lamin* (human lamin C amino acids 66-230), pGBKT7- *p53* (mouse p53 amino acids 72-390), and pGADT7-*Large-T* (SV40 large-T antigen amino acids 84-708) as control plasmids. Mouse *Nxf1* (C57BL/6J) open reading frame and a site-directed E610G mutation (to model CAST/Ei) were cloned into pGBKT7 after high-fidelity PCR as bait proteins. Mouse *Nup62* open reading frame was cloned into pGADT7 as a prey for Nxf1 fusion protein. A 430-bp *CUP1* promoter sequence [23] was synthesized as a gBlock Gene Fragment (IDT), with nucleotide changes to destroy the *Bam*HI and *Nde*I sites, and amplified by Phusion PCR. The pGBKT7 vector was digested to remove the 705 bp *ADH1* promoter, gel purified, and used to clone the modified *CUP1* promoter to create pGBK-*CUP1*. (Figure 1A). Fluorescent proteins EGFP, mCherry, and mGrape3 [27–29] and allelic variants of *Nxf1* were directionally cloned behind the *CUP1* promoter in this new vector at the BamHI and NdeI sites within the multiple cloning site derived from pGBKT7. Plasmids have been deposited with Addgene, including pGBK-*CUP1* empty vector (Addgene 169710) and pGBK-*CUP1* with open reading frames for EGFP (170210), mCherry (170264), mGrape3 (170265), Nxf1-B (170266), and Nxf1-C (170267).

**Figure 1.**
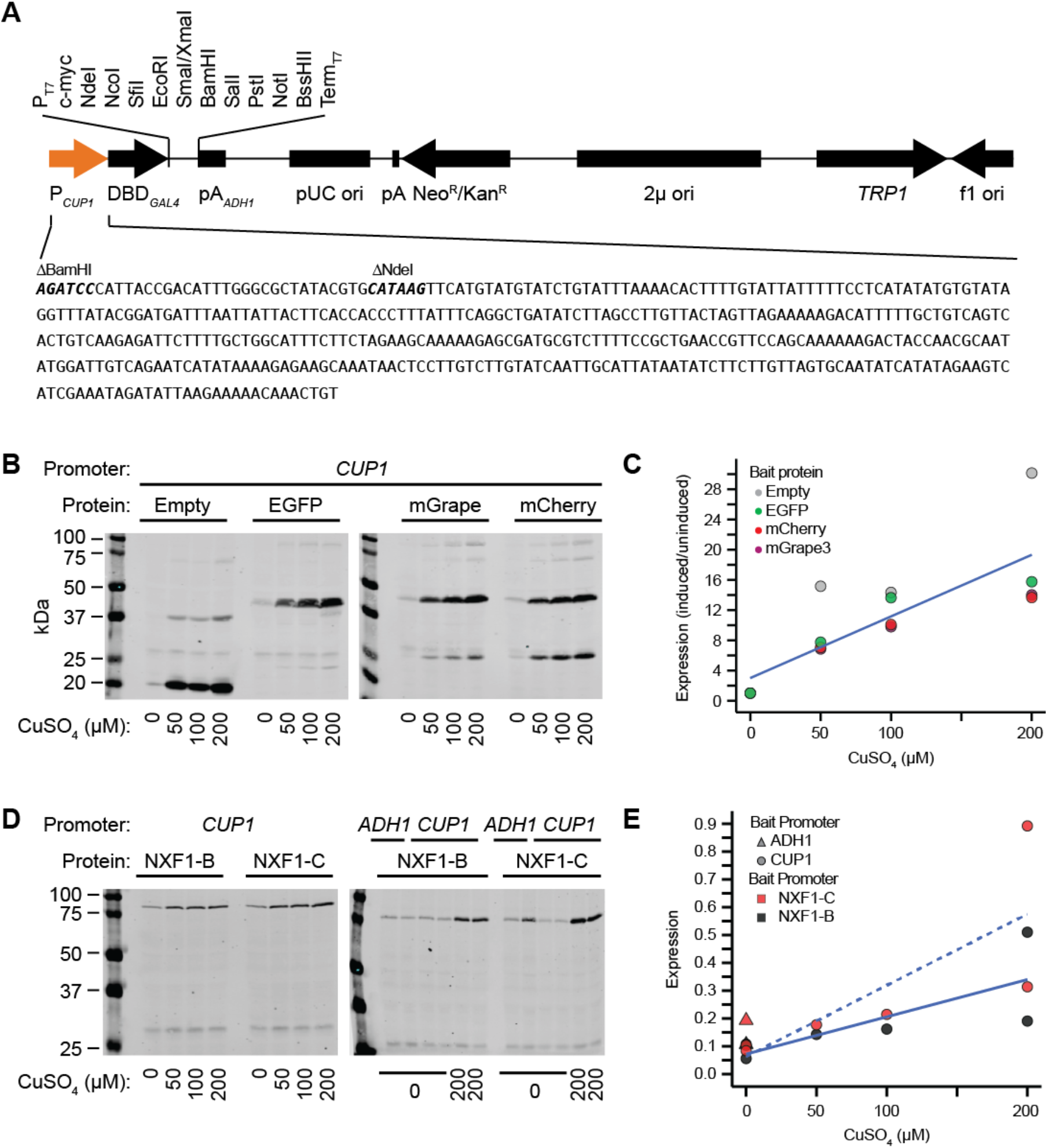
*CUP1* promoter is dosage sensitive to copper ion concentrations. **(A)** Map of the 7-kb modified pGBKT7 vector with *CUP1* promoter (copper arrow). Modified *CUP1* promoter sequence to remove restriction sites is shown below. **(B)** The *CUP1* promoter increased expression in response to an increase in CuSO_4_ concentration. Western blot data showed a change in protein expression as CuSO_4_ concentrations increased from 0 (uninduced) to 50, 100, and 200 μM (induced). Normalized expression levels were plotted relative to 0mM CuSO_4_ (uninduced) and a best fit line was overlayed. **(C)** Western blot data showed greater expression when using an induced *CUP1* promoter compared to the *ADH1* promoter, and greater expression of NXF1-B than NXF1-C. Normalized expression levels were plotted as raw values and a best fit line was plotted only for CUP1 driven NXF1-B and NXF1-C (NXF1-B, *NXF1*^*B*6^; NXF1-C, *NXF1*^CAST^).

### Yeast strains

Yeast transformation and growth were carried out in AH109 (*MATa, trp1-901, leu2-3, 112, ura3-52, his3-200, gal4Δ, gal80Δ, LYS2 :: GAL1_UAS_-GAL1_TATA_-HIS3, MEL1 GAL2_UAS_-GAL2_TATA_-ADE2, URA3::MEL1_UAS_-MEL1_TATA_-lacZ*), obtained as a part of the Yeast Matchmaker Two-Hybrid Systems. Transformation-competent AH109 yeast cells were prepared essentially as described [30]; briefly, cells were grown in 2x YPD media (Fisher Scientific, DF0427-17-6) supplemented with 80 mg/L of adenine hemisulfate, harvested, and stored frozen at −80°C. AH109 was transformed with 200 ng of plasmid for single-transformants and 200 ng of each plasmid for co-transformants using the LiAc/SS carrier DNA/PEG method [30]. Transformed cells were grown on plates containing 1.8% agarose, minimal SD base media (Takara Bio, 630411), and a composition of every essential amino acid except tryptophan (SD/-Trp) (630413), leucine (SD/-Leu) (630414), or tryptophan and leucine (SD/-Trp/-Leu) (630417) to select for auxotrophic markers on either or both plasmids at 30°C for 3-5 days.

### Western blot assay

Single yeast colonies were picked into 5 ml of selective media for liquid culture and incubated ~18-22 hours at 30°C and 220 rpm to reach saturation. Saturated cultures were diluted into fresh selective medium to OD_600_ = ~0.2-0.3 in 7 ml total and incubated ~3 hours at 30°C, 220 rpm, to obtain mid-log phase of OD_600_ = ~0.45-0.65. To induce the *CUP1* promoter, CuSO_4_·5H_2_O (Ricca Chemical, 2330-16) was added prior to the second incubation. After incubation, 5 ml of chilled liquid culture was transferred to pre-chilled tubes. Cells were pelleted at 1000 x g and 4-10°C for 5 minute and washed twice with ice-cold H_2_O. Pellets were stored at −80°C until used for protein extraction. Protein was extracted essentially as described [31]; briefly, cells were resuspended in 0.2M NaOH and incubated for 5 minutes at room temperature, pelleted, and resuspended in SDS sample buffer (0.06M Tris-HCl, pH 6.8, 5% glycerol, 2% SDS, 4% *β*-mercaptoethanol, 0.0025% bromophenol blue) supplemented with protease inhibitor (1% Millipore Sigma P8340, 1% PMSF), incubated at 100°C for 3 minutes, and centrifuged to pellet debris with the lysate retained. The volume of NaOH and SDS sample buffer used was dependent on the final OD_600_ measurement before harvesting to account for variability in growth. Protein lysates were stored at −20°C until used in western blots. Proteins (7 μl of extract) were separated on Laemmli SDS-PAGE gels and transferred to nitrocellulose membranes (Bio-Rad, 1620112). Membranes were stained with Ponceau-S for visualization. Epitope-tagged bait proteins were detected with mouse anti-c-myc monoclonal antibody (Invitrogen, MA1-980) and IR-680 conjugated donkey anti-mouse secondary antibody (LI-COR, 926-68072). Blots were imaged using a LI-COR Odyssey CLx imaging system. As a proxy for total protein, blots were re-probed with an anti-phosphoprotein antibody cocktail (Millipore Sigma, P3430 and P3300) and IR-800 conjugated secondary antibody. Blot images were quantified using ImageJ and data points were corrected using protein signals normalized to total protein.

### Quantitative liquid growth yeast two-hybrid assay

A single co-transformed colony (<2 months old) containing both bait and prey plasmids was picked into 2-3 ml of SD/-Trp/-Leu liquid media. Cultures were incubated overnight for approximately 18-22 hours at 30°C and 220rpm until saturation. The overnight cultures were diluted into fresh SD/-Trp/-Leu media to a constant optical density, then split evenly into SD/-Trp/-Leu/-His media to create an OD_600_ = ~0.13. CuSO_4_·5H_2_O was added to the yeast in SD/-Trp/-Leu/-His media to the specified concentrations. 950 μl of sample was then added to a 24-well flat bottom plate following a matrix format. The plate was incubated at 30°C with rocking in a chamber humidified with the same liquid media to offset evaporation. The samples were resuspended before measuring the OD_612_ every hour. Optical densities were measured on a Tecan i-control Infinite 200 plate reader.

## RESULTS

### A modified Y2H bait plasmid with copper-induced expression

The classical Y2H GAL4 system relies on a 700 bp *ADH1* promoter intended for high, constitutive expression [32]. We replaced the *ADH1* promoter in pGBKT7 with a modified *CUP1* promoter, synthesized to destroy BamHI and NdeI restriction sites so that they remain unique in the multiple cloning site of the resulting plasmid (Figure 1A). To determine whether the modified promoter sequence retained both basal (without exogenous copper) and copper-induced activity, we cloned fluorescent protein open reading frames in-frame with the c-myc epitope tag in the vector and monitored expression of the epitope by infra-red Western blotting (Figure 1B). This showed robust and concentration-dependent induction by copper concentrations 50-200 μM (Figure 1C and S1_Table for empty vector (expressing GAL4 DNA-binding domain and c-myc epitope) and for three out of three fluorescent proteins (EGFP, mGrape3, and mCherry). To validate induction with a more challenging bait protein, we cloned full-length open reading frames for alternate alleles of *Nxf1* into the new vector and compared basal and induced expression to that from the *ADH1* promoter of the original pGBKT7 plasmid (Figure 1D). This again showed concentration-dependent induction of bait protein expression (Figure 1E and S2_Table). Basal (un-induced) expression from the *CUP1* promoter remained detectable, but at lower levels than from the constitutive *ADH1* promoter, while induced expression from *CUP1* achieved a higher level than *ADH1*.

### Y2H liquid growth assay validation for Nxf1:Nup62 interaction

To establish that copper-induced bait expression is compatible with Y2H detection and relative quantification of PPI strength, we implemented a liquid growth assay (Figure 2A). We used a 24-well format with replicate cultures for allelic Nxf1 bait proteins and a known C-terminal interaction partner, Nup62 [33–40], and for positive and negative controls with plate positions alternating across replicates. We measured growth rate by optical density as a function of time while maintaining auxotrophic selection for each Y2H plasmid and the integrated *GAL1_UAS_-GAL1_TATA_-HIS3* reporter in AH109, which requires PPI for growth in the absence of histidine. Growth was evident for both Nxf1 bait proteins with Nup62 prey, with faster growth for the P53 bait and SV40-LargeT prey as strong positive control and essentially zero growth among negative controls (Figure 2A and S3_Table).

**Figure 2.**
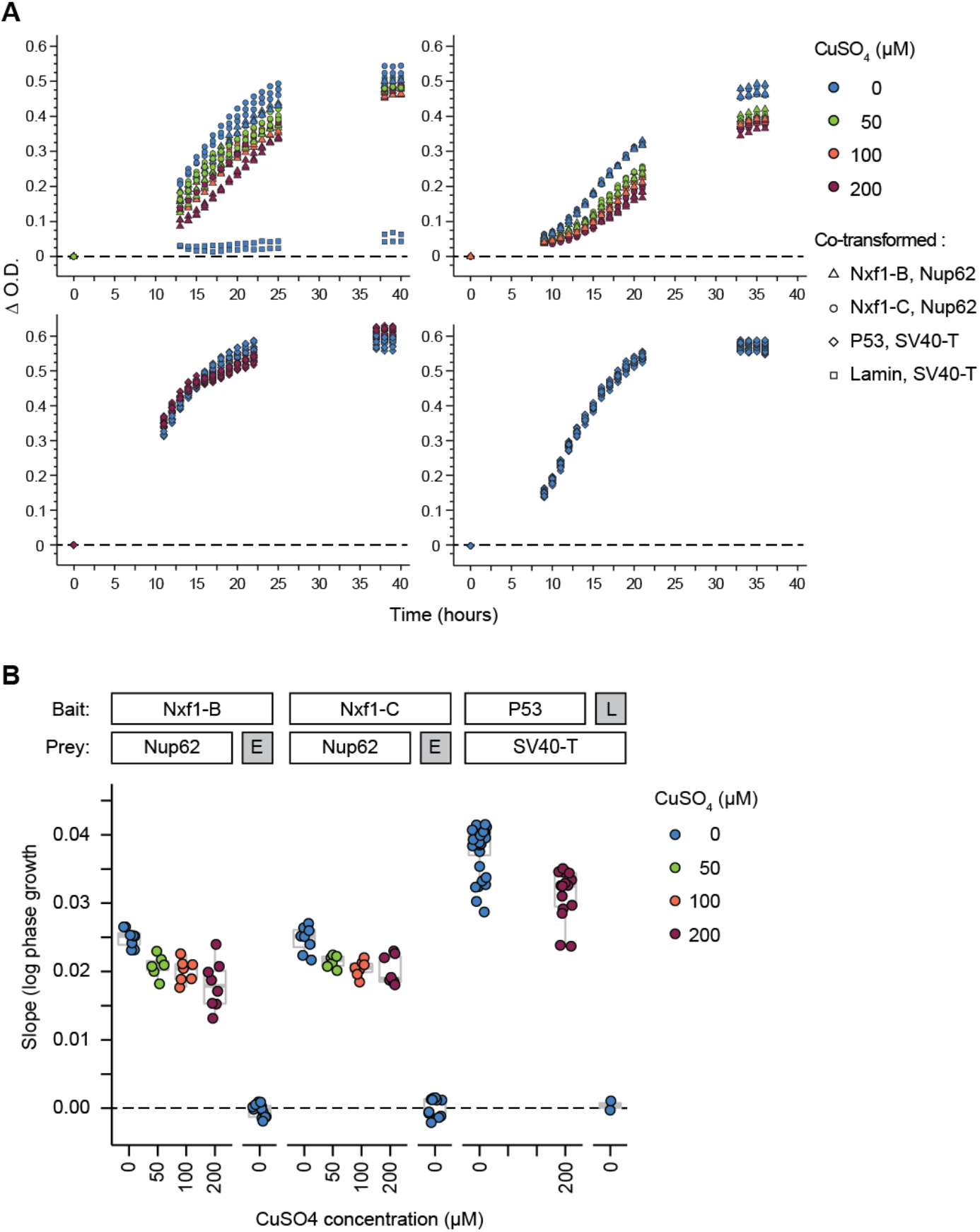
Y2H growth rate decreases at higher copper concentration. **(A)** Representative growth curves of liquid Y2H assays over time. Yeast co-transformed with either allele of *Nxf1* as bait and *Nup62* or empty vector prey, *P53* and *SV40-T* (positive interaction control), or *Lamin* and *SV40-T* (interaction negative control) were cultured in SD/-Trp/-Leu/-His media to select for interaction in a 24-well plate. Samples were grown at 30°C with rocking and measured at OD_612_ in hourly intervals up to 40 hours (NXF1-C, *NXF1*^CAST^; NXF1-B, *NXF1*^*B*6^). Y-axis is change in OD_612_ relative to time 0. **(B)** Quantitative measurements of the growth rate derived from growth curves. The slope during the log linear growth phase was utilized to calculate the growth rate of each sample. Nxf1 bait expressed from with empty prey vector (E) and Lamin bait (L) with SV40 Large T antigen (SV40-T) were negative controls.

We took the slope of the optical density curve during log phase as a measure to visualize growth effects of a given bait-prey pair and copper concentration (Figure 2B). For each bait-prey combination, including the *ADH1*-driven P53 bait, increasing copper concentration decreased growth rate, consistent with its known toxicity in *Saccharomyces* [25, 26] (Figure 2B and S4_Table). Estimates of interaction strength in this assay will thus require controls done at the same copper concentration, at least when using growth rate as the outcome measure.

## DISCUSSION

Yeast two-hybrid approaches have provided a flexible platform for exploring protein interactions for more than 30 years. Here, we introduced a copper-inducible bait vector based on a modified *CUP1* promoter and showed Y2H detection of a known interaction under both low basal expression and highly induced expression of the bait. Adding copper to liquid media allowed high level of bait protein induction and was compatible with detecting PPI, but copper toxicity decreased growth rate independent of induction, as seen by reduced growth for *ADH1*-driven positive control pair in Figure 2. Performance of the *CUP1* Y2H vector might be improved in future studies by transient exposure to copper, or by use of a fluorescent or enzymatic reporter in place of growth rate as an assay. Inducible promoters from other systems or use of synthetic regulators may allow tighter regulation or induction mechanisms that are orthogonal to the biology of *Saccharomyces* to reduce confounding between expression level and growth. The *CUP1* Y2H plasmids described here have been deposited with Addgene.

Our results also showed measurable interaction between NXF1 and NUP62 independent of *Nxf1* (E610G) allele. While growth rates across different copper concentrations should be interpreted with caution, *HIS3* complementation from the Y2H reporter is clear across all copper concentrations and show similar quantitative levels between *Nxf1* alleles for each copper concentration tested. While quantitative growth rate for the same interaction varies across basal expression and copper concentrations for acute induction (or for the un-induced positive control), in each condition tested the permissive (B6) and suppressing (CAST) NXF1 proteins were not significantly different in this assay, suggesting that differential interaction with the FG-repeat containing nucleoporins may not be relevant to *Nxf1*-mediated genetic suppression. Future studies will be required to test whether other known NXF1-interacting proteins might show differences in interaction by the Y2H assay. The approach and data here provide validation for a modified Y2H bait vector to facilitate such studies.

## Supporting information

Data for Figure 1C

Data for Figure 1E

Data for Figure 2A

Data for Figure 2B

## ACKNOWLEDGEMENTS

We thank Professor Peter Novick for suggesting *CUP1* as an inducible system and Professors Nathan Shaner and Roger Tsien for sharing cloned fluorescent protein genes. This work was supported by a grant from the National Institute for General Medical Sciences, R01 GM086912.

